# Chromosomal Fusions and Evolutionary Forces: Exploring the Neo-Sex Chromosome System of *Anolis distichus*

**DOI:** 10.1101/2025.08.05.668756

**Authors:** Cleo H. Falvey, Pietro de Mello, Jody M. Taft, Alyssa A. Vanerelli, Paul M. Hime, Alana M. Alexander, Richard E. Glor, Anthony J. Geneva

**Author notes:** These authors contributed equally to this work as the co-primary and co-senior authors, respectively.

## Abstract

The evolutionary dynamics of sex chromosomes differ from autosomes due to their unique pattern of in-heritance and regions of hemizygosity in non-recombining areas. However, the study of sex chromosomes and sex-linked gene evolution has been limited by the rarity of truly novel sex chromosome complements in model systems. Recent advances in next-generation sequencing have enabled the identification of neo-sex chromosomes, created by the fission or fusion of autosomes with sex chromosomes, providing a new avenue to investigate the dynamics of sex chromosome evolution. Squamate reptiles, particularly *Anolis* lizards, are an excellent system for studying the consequences of sex-linkage due to their frequent sex chromosome-autosome fusions. The Hispaniolan Bark Anole, *Anolis distichus*, has experienced two sex chromosome and autosome fusions that led to a multiple sex chromosome system (X_1_X_2_Y). We present a high-quality whole-genome assembly and annotation of a male *A. distichus* (X_1_X_2_Y), enabling a detailed analysis of all three of its neo-sex chromosomes. We identify scaffolds associated with X_1_, X_2_, and Y chromosomes using an integrative approach and estimate degeneration and selection strength. Our results support long-held theories of differential evolutionary pressures in sex chromosomes, such as the Fast X effect and Y degeneration. Additionally, we find that chromosome 12 has become sex-linked independently in two different *Anolis* species, suggesting that some autosomes may be more likely to become sex-linked. Altogether, our genome adds to the diversity of available taxa sequenced and enables novel comparative analyses in a variety of fields, including speciation, chromosomal synteny, and sex chromosome evolution.

## Introduction

Sex chromosomes play an outsized role in evolutionary processes like adaptation and speciation, and offer the opportunity to study the impacts of reduced recombination and hemizygosity on genome evolution(O’Neill and O’Neill, 2018; Muralidhar and Coop, 2024; Presgraves, 2018; Lund-Hansen et al., 2021; Wilson et al., 2020). In the classical model of sex chromosome evolution, emergence of a new sex-determining locus is quickly followed by evolution of a region with little or no recombination around it. This reduced recombination results in degradation of the chromosome that contains the sex-determining locus, including the accumulation of repetitive elements and loss of functional genes. Muller proposed a ratchet-like process, in which deleterious mutations and transposable elements accumulate but fail to be purged by selection due to strong linkage with the sex-determining region (Muller, 1964; Engelstädter, 2008; Stöck et al., 2021). This degradation is expected to result in heteromorphic sex chromosomes that are clearly distinguishable in traditional karyotypic analyses and regions of the genome that are effectively hemizygous in the heterogametic sex (Kratochvíl et al., 2021; Saunders and Muyle, 2024; Charlesworth et al., 2005; Charlesworth and Charlesworth, 1978).

This hemizygosity has several important consequences: First, it results in smaller effective population sizes for sex-chromosome associated loci (Abbott et al., 2017; Furman et al., 2020) , which further limits the effectiveness of selection and accelerates degradation: Although natural selection is generally less effective in regions of the genome with smaller effective population sizes, hemizygous regions of sex chromosomes often include some of the fastest-evolving genes in the genome, linked to the expression of primary sex traits, also known as the Fast-X effect (Charlesworth et al., 2018; Meisel and Connallon, 2013). This phenomenon is hypothesized to be a consequence of hemizygous genes being exposed to selection as haploids in the heterogametic sex, exposing advantageous recessive alleles to selection more promptly than in autosomes. Altogether, sex chromosome turnover is a dynamic process that creates a huge diversity of sex chromosome complements, even within closely-related taxa (Palmer et al., 2019; Rovatsos et al., 2015).

Because sex chromosome degradation is expected to occur over thousands, or even millions of generations, studying the genomic and evolutionary consequences of sex chromosome evolution requires systems with sex chromosomes that are experiencing varying degrees of degeneration. Unfortunately, most model organisms either have extensively degenerated sex chromosomes that are easily identifiable as heteromorphic in traditional karyotypes (e.g., mice, chicken, fruit flies), or apparently relatively undifferentiated sex chromosomes that are homomorphic in a karyotype (e.g., African clawed frogs) (Cortez et al., 2014; Zhou et al., 2014; Miura, 2017; Tymowska and Kobel, 2008). Organisms possessing sex chromosomes that are the result of relatively recent rearrangements of ancestral sex chromosomes and autosomes can provide unique opportunities to study sex chromosome evolution (Kitano et al., 2009; Lisachov et al., 2023; Mongue et al., 2017). Such species allow for the direct comparison of (older) degenerated hemizygous regions and (younger) recently sex-linked regions, investigation of the dynamics of chromosomal fusions and fissions, and provide insight into the effects of these chromosomal rearrangements on broad evolutionary patterns (Lenormand et al., 2020; Zhu et al., 2022; Filatov, 2018).

Although prior work suggests that both X and Y chromosomes are capable of fusing with autosomes, Y-autosome fusions are more common, typically resulting in an XXY system: A neo-Y, formed by fusion of the ancestral Y and an ancestral autosome, is homologous with two distinct Xs - the ancestral X and a neo-X representing the partner of the autosomal chromosome that fused to the ancestral Y (Pennell et al., 2015). Advances in comparative genomic analyses have allowed for the identification of the chromosomes involved in these fusions and have revealed another potential general feature of sex chromosome evolution – namely, the tendency for the same specific ancestral autosomes to be repeatedly co-opted into becoming sex chromosomes (Kichigin et al., 2016; Giovannotti et al., 2017; Lisachov et al., 2019; O’Meally et al., 2012; Sardell et al., 2021).

Squamate reptiles (lizards, snakes, and amphisbaenians) are ideal models for studying sex chromosome evolution due to their diverse sex chromosome complements, which are often the result of sex chromosome and autosome rearrangements (Mezzasalma et al., 2021; Pennell et al., 2015; Pinto et al., 2023, 2022). This is especially true for *Anolis* lizards, which have long served as a model system in ecological and evolutionary research and have now become a model for genomic studies, as they were the first lizard species to have a fully sequenced genome (Alföldi et al., 2011; Losos, 2009). Although the large clade of iguanian lizards to which *Anolis* belongs is believed to have genetic sex determination, understanding sex chromosome evolution in this group has long been complicated by the fact that many species are karyotypically homomorphic. However, genome sequencing of the green anole (*Anolis carolinensis*) – the first non-avian reptile genome sequenced –resulted in the identification of its putative X chromosome (Alföldi et al., 2011). Subsequent cytogenetic and phylogenetic comparative analyses showed *Anolis carolinensis* had an XY sex chromosome system, and that the X chromosome is likely ancestral across iguanians (Gamble et al., 2014; Rovatsos et al., 2014a; Lisachov et al., 2019, 2017; Alföldi et al., 2011; Pokorńa and Kratovchíl, 2009). Additionally, fusions and fissions involving ancestral sex chromosomes and autosomes are common across anoles, resulting in a diversity of sex chromosome complements (Gamble et al., 2014; Gorman and Atkins, 1969; Castiglia et al., 2013; Webster et al., 1972; Blake, 1986; Altmanová et al., 2018; Rovatsos et al., 2014c; Pennell et al., 2015; Rovatsos et al., 2014a).

*Anolis distichus*, native to the Bahamas and Hispaniola, possesses an X_1_X_2_Y sex chromosome complement, previously identified via cytogenetics, flow-sorted chromosome sequencing, and fluorescence in-situ hybridization (Gorman and Atkins, 1966; Gorman, 1974; Guyer and Savage, 1992; Gamble et al., 2014; Kichigin et al., 2016). Previous work comparing *A. distichus* to *A. carolinensis* indicates that this X_1_X_2_Y system likely resulted from two sex chromosome-autosome fusions. One fusion likely occurred between the ancestral anole Y and a microchromosome, resulting in a neo-Y and its homologous X_1_ and X_2_ chromosomes. The other fusion likely occurred between the ancestral anole X and another microchromosome (Lisachov et al., 2019; Giovannotti et al., 2017; Kichigin et al., 2016) (Fig. 1). The order of these events is still unclear, but the existence of two neo-X chromosomes allows us to study sex chromosome degeneration by comparing the extent of divergence of genes in each neo-sex linked region to the ancestral X chromosome. Furthermore, these two fusions have generated neo-X and neo-Y chromosomes, providing a unique opportunity to study the early stages of sex chromosome evolution in both X and Y chromosomes.

**Figure 1:**
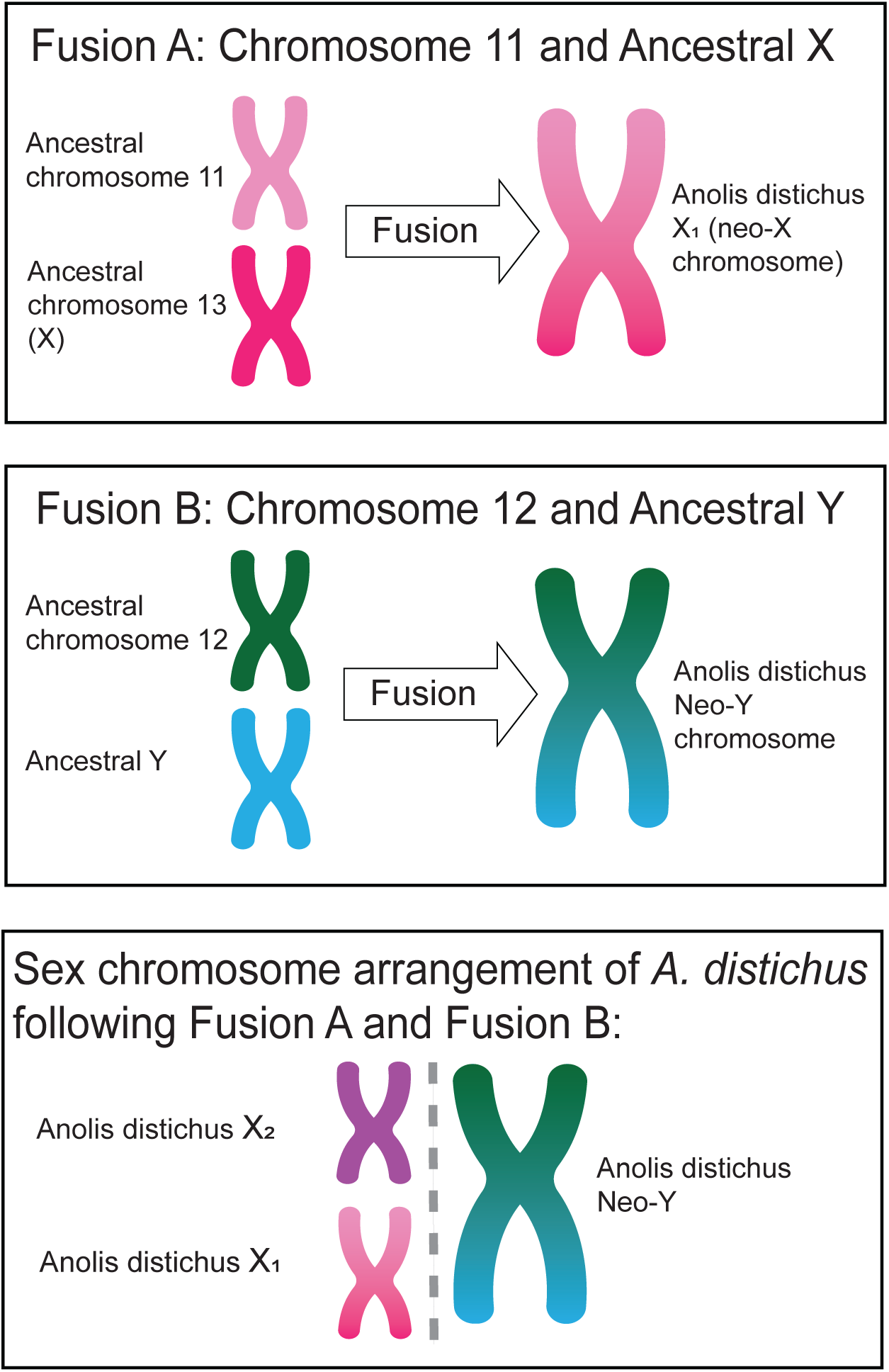
Hypothesized chromosomal rearrangements in *A. distichus*. Schematic showing the hypothesized chromosomal rearrangements between the ancestral X in *A. carolinensis* to the derived karyotype of *A. distichus* based on previous flow-sorting sequencing data (Kichigin et al., 2016; Lisachov et al., 2019; Giovannotti et al., 2017). **Fusion A:** In one fusion, the ancestral *Anolis* X chromosome (chromosome 13 in *A. carolinensis*) fused with an ancestral autosome (microchromosome 11 in *A. carolinensis*) resulting in the neo-X_1_ chromosome in *A. distichus*. **Fusion B:** In the second fusion, the ancestral *Anolis* Y chromosome fused with an ancestral autosome (microchromosome 12 in *A. carolinensis*), resulting in the neo-Y chromosome in *A. distichus*. With a lack of intermediate data between the ancestral and derived karyotype, it is not possible to identify the order of the fusions, so they have been referred to as A and B for the purposes of this figure.

In this study, we had 3 main goals: (i) We identified the sex chromosomes of *Anolis distichus* and tested for the presence of previously-hypothesized sex chromosomes and autosome fusions using comparative synteny methods (Giovannotti et al., 2017; Lisachov et al., 2019; Kichigin et al., 2016). (ii) Once identified, we hypothesized that neo-sex chromosomes in *A. distichus* would experience evolutionary pressures consistent with predicted theory (Wright et al., 2016; Furman et al., 2020; Palmer et al., 2019): Specifically, the neo-Y should experience degeneration with gene loss and accumulation of transposable elements (Bachtrog, 2013; Charlesworth and Charlesworth, 2000; Mank, 2012), and the neo-X should experience the Fast X effect (Meisel and Connallon, 2013; Charlesworth et al., 2018). (iii) We predicted that the autosomes involved in these fusions are non-random, with some more likely to be repeatedly recruited to becoming sex-linked than others.(O’Meally et al., 2012; Sigeman et al., 2019; Sardell et al., 2021) To accomplish these goals, we generated and analyzed a new, high quality genome assembly and annotation for *A. distichus*. Altogether, this assembly, annotation, and associated analyses present a comprehensive approach to identify the broadscale consequences of neo-sex chromosome fusions, and enables future investigation in to the effects of these fusions in diversification and adaptation.

## Methods

### Sample Collection

#### Sample Collection

We obtained DNA from tissues sampled from a single adult male *Anolis distichus*. The individual we sequenced resulted from captive breeding of two wild-caught individuals sampled from the Dominican Republic along an unpaved road that extends north from the town of Baní along the Rio Baní (18.3259*^◦^* N, 70.3464*^◦^* W). All protocols for captive lizard husbandry (De Meyer et al., 2019) and sampling were approved by the University of Rochester’s Animal Care and Use Committee (UCAR-2007-030R) and the University of Kansas’s Institutional Animal Care and Use Committee (AUS-227-01).

### Sequencing

#### Pacific Biosciences HiFi Sequencing

We contracted the University of Delaware Sequencing and Genotyping Center to perform Pacific Biosciences (PacBio) high-fidelity long-read sequencing. We extracted high molecular weight (HMW) genomic DNA using the MagAttract HMW DNA Kit (Qiagen Inc., Venlo, Netherlands). We confirmed the presence of HMW DNA in the resulting extractions using pulse-field capillary electrophoresis with a FemtoPulse (Agilent, Santa Clara, CA). We converted 10 *µ*g of extracted DNA into SMRTBell templates using the HiFi Express Template Prep Kit 2.0 (Pacific Biosciences, Menlo Park, CA). We cleaned the resulting templates by removing unligated adapters and damaged DNA fragments via exonuclease treatment, and removed small fragments (*<*5kbp) via pulse-field electrophoresis on a BluePippin (Sage Science Inc., Beverly, MA). We then used 1 *µ*L from the cleaned library as input for a second FemtoPulse analysis that verified size distribution and DNA concentration (Advanced Analytical Technologies Inc., Ankeny, IA). Following verification, we annealed primers and bound DNA polymerase to the two libraries using the Sequel II Binding Kit 2.0 (Pacific Biosciences, Menlo Park, CA). We sequenced the resulting libraries on the PacBio Sequel IIe with 8M SMRT cells using chemistry 3.0, with four hour pre-extensions and 30 hour movie time. The 49 million PacBio HiFi reads generated by the protocol produced 38.1*×* coverage (Table 1).

**Table 1:** Sequencing effort: Values showing the sequencing effort completed for the *AnoDis1.0* assembly constructed from PacBio HiFi reads and Dovetail Omni-C reads, including the number of bases and reads sequenced, as well as depth, breadth, and proportion of the reads mapping back to the reference.

|  | Pacific Biosciences High Fidelity Reads | Dovetail Omni - C Reads |
| --- | --- | --- |
| Number of Bases Sequenced | 67296365909 | 123418965600 |
| Number of Reads | 24252093 and 24478209<br>(48730302 total) | 411396552 (411360559 after filtering) |
| Average Read Depth | 38.1X | 48.4X |
| Breadth of Coverage | 99.9997% | 99.9829% |
| Reads Mapped to Reference | 98.64% | 99.33% |

#### OMNI-C Sequencing

We contracted Dovetail Genomics to generated Omni-C libraries. Their protocol fixes intact chromatin with formaldehyde prior to using proximity ligation to preserve the 3D conformation of nuclear DNA. Dovetail Genomics then created high throughput NEBNext Ultra sequencing libraries using Illumina (Illumina, San Diego, CA) adapters from purified, ligated DNA fragments (New England Biosciences, Ipswich, MA). Sequencing of our OMNI-C libraries on the Illumina HiSeqX platform resulted in 41 million reads and 48.4*×* coverage (Table 1.)

### Genome Assembly and Annotation

#### Assembly Process

We performed our initial genome assembly with PacBio long read data using Hifiasm v.0.16.1, which assembled each haplotype before creating a primary assembly (Cheng et al., 2021). We then scaffolded this assembly using YAHS v1.2a.2 (Zhou et al., 2022), which mapped the Omni-C reads to the Hifiasm assembly using the Arima Genomics pipeline (https://github.com/ArimaGenomics/mapping_ pipeline). We ran BUSCO v5.2.2 with the vertebrata odb10.2019-11-20 dataset (Simão et al., 2015) and the stats function from BBMap v.38.97 (Bushnell, 2014) before and after scaffolding to ensure that the completeness and contiguity improved iteratively. To identify potential errors, we also aligned the Omni-C reads back to the scaffolded assembly using Juicer v.1.6, and visualized the Omni-C contact maps created using JuiceBox v2.3.0 (Durand et al., 2016b; Robinson et al., 2018) (Fig. 2). Our final assembly’s genome size of 2.08 Gbp was within the expected range based on previously published anole genomes, which range from 1.78 Gbp (Alföldi et al., 2011) to 2.4 Gbp (Pirani et al., 2023).

**Figure 2:**
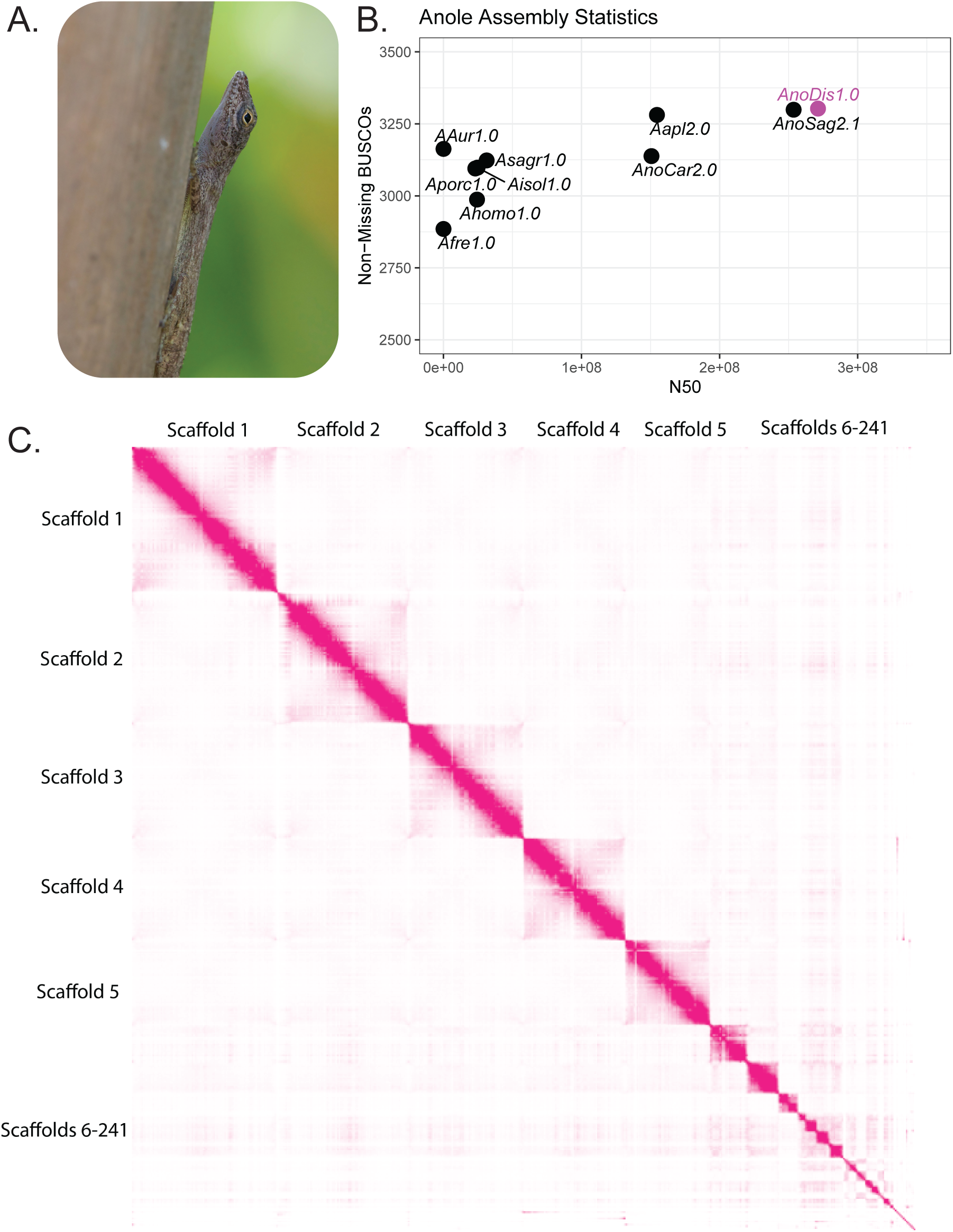
*AnoDis1.0* assembly information: **A.** *Anolis distichus*, photo by Richard Glor. **B.** Assembly statistics (N50 and Non-Missing BUSCOs) of previously assembled *Anolis* lizards (black). The *AnoDis1.0* assembly is highlighted (pink) and is on par with other reptile genomes (such as *A. sagrei)* that have been assembled from PacBio HiFi reads (Geneva et al., 2022). **C.** Link-density histogram results, visualized using JuiceBox tools, labeled with their respective scaffolds on the axes (Durand et al., 2016a,b). Pink regions indicate more than 2060 reads mapping and white regions indicate little to no read pairs mapping in that

#### Annotation Analyses

To obtain transcriptomic data for genome annotation, we used RNAseq data from four *Anolis distichus* tissue samples; three previously published datasets from different colored skin fragments (orange, yellow and white) (de Mello *et al.,* 2021) and a fourth new transcriptome from a 20-30 mg section of non-regenerated tail tip sampled from of a single male *A. distichus* collected from the same locality as the specimen used in the genome assembly. The new transcriptome was processed as detailed in (De Mello et al., 2021).

We contracted DoveTail Genomics (Dovetail Genomics, Scotts Valley, CA) to perform genome annotation. To identify repetitive regions of the genome, Dovetail used RECON v.1.08 (Bao and Eddy, 2002), RepeatScout v1.0.6 (Price et al., 2005), and RepeatModeler v.2.0.1 (Flynn et al., 2020). We then used the results as input for RepeatMasker v4.1.0. Dovetail trained gene models from previously published coding sequences for other squamates – including two other anole species (*Anolis carolinensis* and *Anolis sagrei*), one additional pleurodont iguanian (*Sceloporus undulatus*), one acrodont iguanian (*Pogona vitticeps*), and a more distantly-related teiid lizard (*Salvator merianae*) – by running two programs in parallel: AUGUSTUS v2.5.5 (Stanke and Morgenstern, 2005), which was run through six iterations, and SNAP v.2006-7-28 (Korf, 2004). Dovetail mapped RNAseq reads onto the genome using STAR v.2.7 (Dobin et al., 2013) and generated intron hints with the bam2hints command in AUGUSTUS. Predictions from SNAP and AUGUSTUS were supplemented with the previously published squamate gene datasets and Swiss-Prot peptide sequences from the UniProt database using the MAKER pipeline (Cantarel et al., 2008). Dovetail predicted genomic regions that encode tRNA using tRNAscan-SE v2.0.5 (Chan and Lowe, 2019). To assess the quality of gene predictions, we used the MAKER pipeline to generate annotation edit distance (AED) scores for each putative coding sequence (Cantarel et al., 2008). To generate final gene models, we conducted two filtering steps. In the first step, we removed regions that were not predicted by both MAKER and AUGUSTUS, or that did not have a positive BLAST search result when queried against the UniProt database. In the second step, we removed all remaining unidentified genes that did not contain three or more exons.

### Identifying Sex Chromosomes

Prior work hypothesizes that *A. distichus* possesses two X chromosomes: one of these, (X_1_), was presumably derived from a fusion between the ancestral X chromosome (scaffold 13 in *A. carolinensis*) and an ancestral autosome (microchromosome 11 in *A. carolinensis*) (Lisachov et al., 2019; Giovannotti et al., 2017; Kichigin et al., 2016) (Fig. 1). The other X chromosome (X_2_), presumably resulted from the fusion of an ancestral autosome (likely microchromosome 12 in *A. carolinensis*) and the ancestral Y chromosome, establishing an X_1_X_2_Y system (Fig 1).

To test the hypothesis that the XXY genome of *A. distichus* resulted from these two different fusions, each involving an ancestral sex chromosomes and an ancestral autosome, we identified candidate sex-linked regions within *AnoDis1.0* using two approaches. First, given that our *A. distichus* genome was obtained from a single male specimen, we predicted that both X and Y-linked scaffolds would exhibit half the depth of coverage of autosomes. We tested this prediction by mapping reads the Omni-C reads back to the assembly and looking for scaffolds with significantly lower coverage depths. Second, we predicted that the majority of previously identified genes located on the X and Y chromosomes of *A. carolinensis* should be present in our candidate sex-linked scaffolds, because this species is presumed to have the ancestral X and Y anole chromosomes (Gamble et al., 2014). To test this prediction, we implemented a BLAST search of X- and Y-linked genes identified in *A. carolinensis* against the *A. distichus* genome (Geneva et al., 2022; Marin et al., 2017; Rupp et al., 2016; Rovatsos et al., 2014b).

#### Comparative analyses of sex chromosome synteny

After identifying candidate sex-linked scaffolds based on read mapping and the presence of shared genes with the ancestral anole sex chromosomes, we conducted genome wide synteny analyses by aligning the *A. distichus* genome to that of *A. carolinensis* (*Anolis* clade - GCA 000090745.2) (Alföldi et al., 2011) using GENESPACEv1.3.11 (Lovell et al., 2022; Poe et al., 2017). We used different synteny analyses for X and Y chromosomes because the ancestral Y chromosome is not well characterized in *A. carolinensis*.

In the case of the X chromosomes, we generally expected that portions of the sex-linked scaffolds in *A. distichus* would show high synteny with sex chromosomes previously identified in these two other anole species. We specifically tested two predictions about X chromosome synteny by comparing *A. distichus* with two other anole species (*A. carolinensis* and *A. sagrei*), based on prior work: (1) the X_1_ chromosome in *A. distichus* would be syntenic with *A. carolinensis* chromosome 13 (the ancestral X) and microchromosome 11 (an ancestral microchromosome) and (2) X_2_ would be syntenic with *A. carolinensis* microchromosome 12 (Giovannotti et al., 2017; Lisachov et al., 2019; Kichigin et al., 2016). We implemented GENESPACE with default settings to determine orthology-based syntenic regions simultaneously across the two other species(Lovell et al., 2022). GENESPACE first aligned amino acid sequences across species using DIAMOND2 (Buchfink et al., 2021) and identified orthogroups using OrthoFinder (Emms and Kelly, 2019). Then, using both graph and cluster-based approaches, it extracted syntenic regions from these data for each non-reciprocal pair of genomes in the dataset. Lastly, it collapsed these syntenic orthogroups in a pan-genome annotation. We used the *plot riparian* function in GENESPACE to produce genome-wide synteny plots across all three species as well as a plot focused exclusively on the sex chromosomes (Fig. 3 and Supplementary Fig. 1).

**Figure 3:**
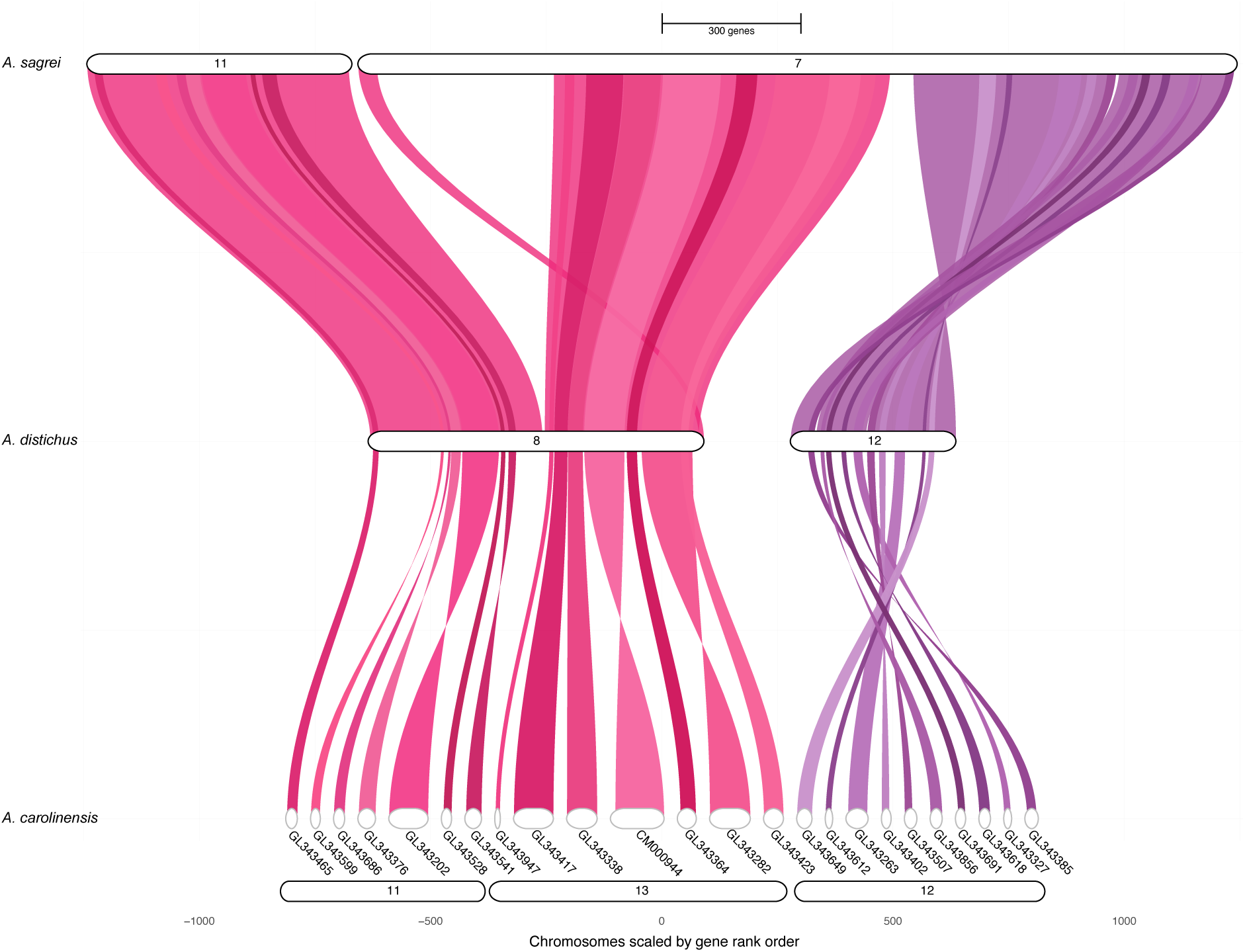
Synteny between X chromosomes in three *Anolis* species: GENESPACE synteny results comparing the sex-linked chromosomes of *A. carolinensis*, *A. sagrei*, and *A. distichus* shows evidence for the sex chromosome and autosome fusions previously identified in *A. distichus*. First, *A. distichus* scaffold 8 (X_1_) results from fusion between *A. carolinensis* chromosomes 11 (autosome) and 13 (X chromosome). Second, *A. distichus* scaffold 12 (X_2_) (the partner of this species’ neo-Y) is syntenic to *A. carolinensis* chromosome 12 and *A. sagrei* chromosome 7 (the neo-X chromosome in *A. sagrei*, which is a result of a fusion between the ancestral X and chromosome 12).

In the case of the Y chromosome, we relied on synteny comparing candidate sex-linked scaffolds from *A. distichus* to one another, rather than assessing synteny across the genomes of anole species. Since the previously autosomal regions of the neo-sex chromosomes are relatively young (roughly 35 million years old compared to an estimates of around 85 million years old for the ancestral X), we predicted that the non-degenerated Y and pseudo-autosomal regions would be syntenic between the candidate X_1_ and neo-Y chromosomes (Román-Palacios et al., 2018; Zheng and Wiens, 2016; Rovatsos et al., 2014c). To test this prediction, we performed a synteny analysis where each of the four candidate sex-linked scaffolds were queried against a database comprised of the remainder of the *A. distichus* genome using SatsumaSynteny v.2.0 (Fig. 4) (Grabherr et al., 2010). SatsumaSynteny identifies syntenic regions through pairwise comparisons by finding long regions (*>*4096 bp) of high similarity. When running *AnoDis1.0* sex-linked scaffolds against themselves, we hard-masked regions annotated as repetitive by converting to Ns. To allow for expected divergence between gametologs, we decreased the minimum probability of a match to 0.999 to qualify for a match (from 0.9999).

**Figure 4:**
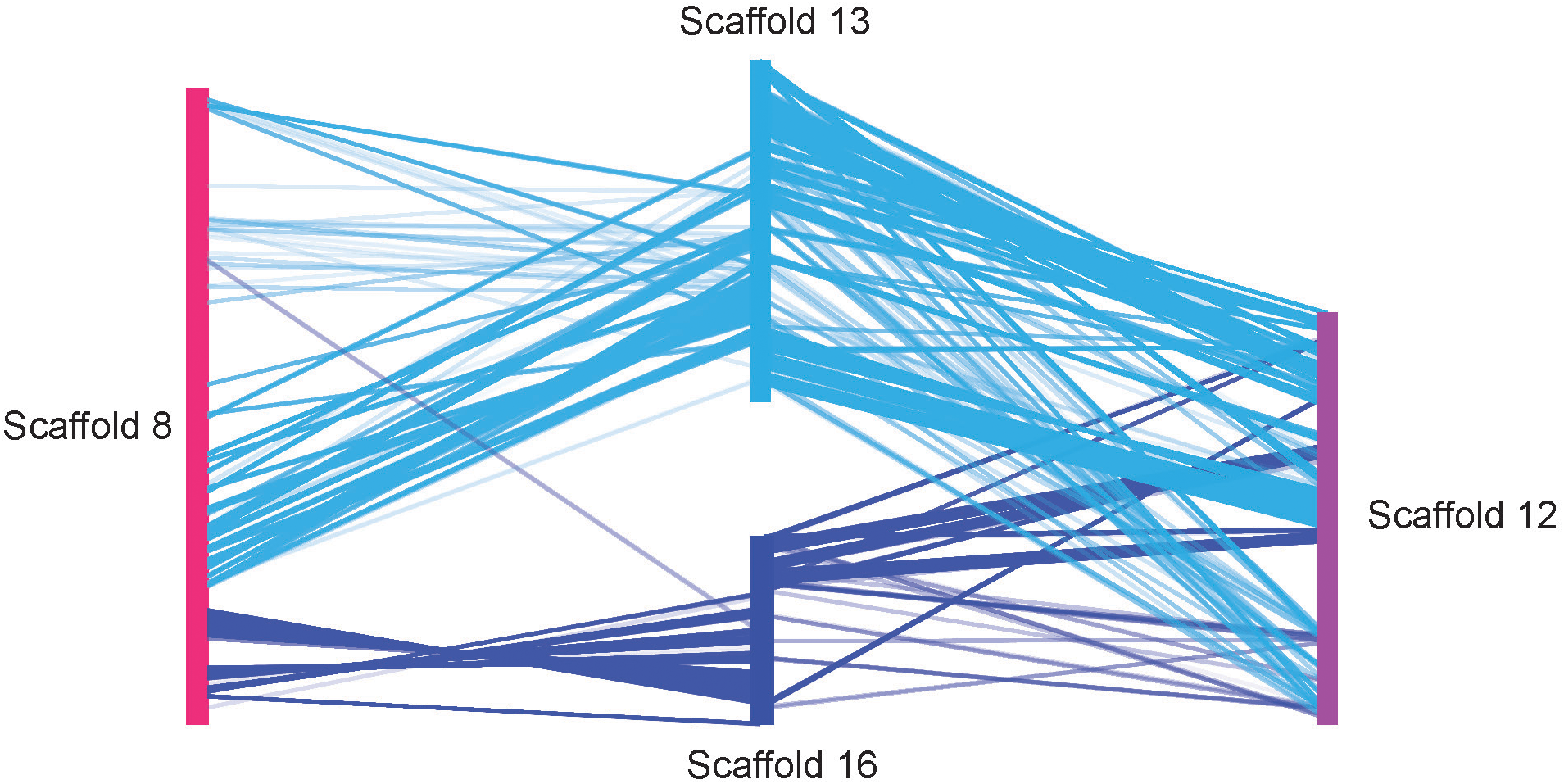
Synteny of *Anolis distichus* mapped to itself, highlighting the relationship between sex chromosomes: Satsuma Synteny results of *A. distichus* sex-linked scaffolds mapped to the remainder of the genome. Large swaths of the hypothesized neo-X_1_ (scaffold 8) and neo-X_2_ (scaffold 12) are syntenic to the neo-Y (scaffolds 13 and 16). Scaffold 8 is largely syntenic to the *A. carolinensis* ancestral X chromosome and contains more than 85% of the known X-linked genes. Combined with the lack of similar levels of synteny between scaffolds 8 and 12, these results strongly support classification of scaffold 8 as X_1_ and scaffold 12 as X_2_.

### Testing Predictions Regarding Sex Chromosome Evolution

In addition to identifying sex chromosomes and the fusions they underwent, we used our genomic sequence data to test two hypotheses about how sex chromosomes are expected to evolve relative to autosomes: (1) First, we predicted that Y-linked regions would have lower gene density, higher transposable element density, and a higher proportion of high and moderate impact SNPs, given the degeneration of gametologs on the Y chromosome (Charlesworth and Charlesworth, 2000; Abbott et al., 2017; Bachtrog, 2013).(2) Additionally, we predicted that both neo-X_1_ and neo-X_2_ should be experiencing evolutionary signatures of the Fast X effect, which would include signatures of positive selection, including a higher proportion of high and moderate effect variants, and higher K_a_/K_s_ values, a proxy for selection strength (Darolti et al., 2023; Li, 1993; Charlesworth et al., 2018). We tested these hypotheses by characterizing the distribution of variant sites and the prevalence of repetitive content across *AnoDis1.0*.

#### Variant Calling

To test our predictions about sex chromosome evolution, we first used the Genome Analysis Toolkit pipeline (Van der Auwera and O’Connor, 2020) (GATK) to call variants within our genome. First, we used BWA-MEM v.0.7.17 (Li and Durbin, 2009) to map the Hi-C reads back to the assembled genome. We then ran HaplotypeCaller in base-pair resolution mode, followed by GenotypeGVCF with the all-sites setting to retain invariant sites (Poplin et al., 2017). Lastly, we filtered likely erroneous variant calls using both GATK v.4.2.2.0 and VCFtools v0.1.16-13 with standard settings (MAPQ *>* 30) and a maximum depth filter of 83.3811 (twice the mean depth) (Danecek et al., 2011). We validated SNP calls by aligning sequencing reads back to the reference, with the expectation that virtually all of the putative variant sites would be heterozygous for both the reference nucleotide and a second allele.

We used our final set of SNPs as input for the program SnpEff, which annotated SNPs and predicted their effects on genes and protein sequences (Cingolani et al., 2012). We created a custom database in SnpEff built from the AnoDis1.0 assembly. SnpEff classified each SNP as high, moderate, or low impact, and identified the impacted region of the genome. We used our SnpEff VCF file to calculate the proportion of high and moderate effect variants and compared the distribution of these proportions between sex chromosomes and autosomes for one megabase windows across the genome, with the expectation that sex chromosomes would have a larger proportion of high and moderate effect SNPs than autosomes as they accumulate deleterious variants. Lastly, we used the *kaks.py* script from SelectionTools v1.1 to calculate the synonymous to nonsynonymous substitution rate for each gene in the assembly, as we hypothesized that sex chromosomes would be experiencing a faster rate of positive selection due to the Fast X effect (Cadzow et al., 2014).

#### Repetitive Element Landscape

We modeled repeats *de novo* using RepeatModeler v2.0.2 (Flynn et al., 2020) and annotated the repeat consensus sequences using RepeatMasker v4.1.2 (Smit et al., 2013-2015). To maximize repetitive element identification, we used four repeat libraries as references: 1) simple repeats, 2) the Tetrapoda DFam library v3.7, 3) previously classified repeat elements from the *A. distichus*-specific *de novo* repeat library generated with RepeatModeler, and 4) unknown repeat elements from the *A. distichus*-specific *de novo* repeat library. We generated alignments for each repeat family and calculated the Kimura-2 parameter divergence from consensus (correcting for CpG sites) using the calcDivergenceFromAlign.pl RepeatMasker tool (Supplemental Figure 2). We used divergence between an insert from its family consensus as a proxy for its age, under the assumption that older elements accumulate more mutations from the consensus. However, we did not convert these divergence values into absolute time estimates, as doing so would require a reliable substitution rate calibrated to known divergence times in *Anolis distichus*. Our use of K2P divergence is thus strictly for comparing relative repeat activity levels across repeat classes, rather than inferring exact ages of insertion.

#### Sliding Window Analyses

We predicted that the Y chromosome would be experiencing degeneration as measured by gene and transposable element density. To compare the distribution of repetitive element and gene densities between sex scaffolds and autosomal scaffolds, we used Bedtools2 to create sliding windows across the genome that were 1 megabase in length (Quinlan and Hall, 2010). We counted how many repeat elements and genes were present in each 1 megabase window and used a Wilcoxon Signed-Rank Test to identify which attributes had a statistically significant difference between autosomes and sex chromosomes. Statistical analyses and visualization were performed in R v4.3.2 using the Tidyverse packages (R Core Team, 2021; Wickham et al., 2019).

### Repeated Recruitment of Sex Chromosomes

Independent sex chromosome and autosome fusions have occurred throughout *Anolis*, and results from other taxa show that some autosomes may be preferentially recruited to become sex chromosomes (Gamble et al., 2014; O’Meally et al., 2012; Sardell et al., 2021; Kitano et al., 2009). To test the hypothesis that certain chromosomes were repeatedly recruited as sex chromosomes, we aligned *A. distichus* to *A. sagrei* (*Norops* clade - GCF 025583915.1) (Geneva et al., 2022) using GENESPACE (Lovell et al., 2022) as outlined above. Previous research shows that *A. sagrei* has a neo-x chromosome as a result of a sex chromosome - autosome fusion, and we predicted that one or more autosomes involved in these fusions were also involved in the sex chromosome fusions in *A. distichus* (Geneva et al., 2022).

## Results

### Genome Assembly and Annotation

#### Assembly Statistics

The assembly has an N50 of 271 MB, and 50% of the assembly is contained in the first four scaffolds (L50=4). The five largest chromosomes were syntenic across all three species (Fig. 1). Additionally, the assembly contained 3188 out of the 3354 (95%) conserved complete single-copy vertebrate genes (Simão et al., 2015; Manni et al., 2021). The original annotation from DoveTail identified 32,350 putative gene models; of those, 13,602 had an identity assigned to them via the Maker pipeline. The remaining 18,748 genes were filtered by the presence of three or more exons, meaning that 7,625 genes with less than three exons were dropped. Our final annotation identified 26,373 genes. The annotated protein set contains 2725 (81.3%) of the conserved, expected vertebrate genes. The total repetitive element content for the *AnoDis1.0* assembly was estimated to be 51.26% of the genome, with 47% comprised of known interspersed repeats and 7.78% unclassified repeats. Retroelements represented 30.52% of total estimated elements (26.27% LINEs, 1.25% SINEs, and 3% LTR elements), while DNA transposons accounted for 8.69% of the detected elements (Supplementary Fig. 2).

### Sex Chromosome Identification

#### Identification of Candidate Sex-Linked Scaffolds

We identified four scaffolds, 8, 12, 13 and 16, as candidate sex chromosomes based on their significantly lower sequencing coverage relative to surrounding similarly-sized autosomes (scaffolds 9, 10, and 11), which all had more than 50X coverage depth (scaffold 8, W=129361, p=2.2 *×* 10*^−^*^16^, scaffold 12 W=152844,p=2.2 *×* 10*^−^*^16^, 13 W=36782, p=2.2 *×* 10*^−^*^16^ and scaffold 16 W=17925, p=2.2 *×* 10*^−^*^16^) (Supplementary Fig. 3).

#### Identifying Y Chromosomes

Direct comparison to the entire ancestral Y chromosome was hindered by the fact that the *A. carolinensis* genome was assembled from a female. However, our BLAST queries found that five out of seven (71.4%) previously identified Y-linked *A. carolinensis* genes mapped to either scaffold 8 or 13 in our *A. distichus* genome, suggesting that scaffold 13 likely consists of part of the ancestral *Anolis* Y chromosome (as scaffold 8 was identified to be neo-X_1_, see below) (Marin et al., 2017). With scaffold 8 putatively assigned to neo-x_1_ and scaffold 13 putatively assigned to neo-X_2_, we predicted that scaffolds 13 and 16, which also had half the depth of coverage of autosomes, might be portions of the Y chromosome. To test whether scaffolds 13 and 16 are parts of the Y chromosome, we performed synteny analyses of all four scaffolds (8, 12, 13, and 16) to the remainder of the *A. distichus* genome (Fig. 4). If scaffolds 13 and 16 are part of the Y chromosome, then scaffolds 8 and 12 should each be syntenic to regions in scaffolds 13 and 16, but scaffold 8 and 12 should not be syntenic to each other. As expected under this scenario, synteny analyses recovered large swaths of scaffold 8 and 12 as syntenic to both scaffolds 13 and 16. Importantly there was no synteny between 8 and 12 (Fig. 4). Additionally, scaffolds 13 and 16 have more than ten times greater number of contacts between each other than between other similarly-sized chromosomes (Supplementary Fig. 5, 4).Therefore, it appears that the Y chromosome was assembled into two pieces: scaffolds 13 and 16.

#### Identifying X Chromosomes

To identify both of the neo-X chromosomes in *A. distichus*, we used a combination of BLAST searches and syntenic comparison to the ancestral *A. carolinensis* X chromosome. First, our BLAST search found that 256 of the 296 genes (Geneva et al., 2022) on the *A. carolinensis* X chromosome had homologs on Scaffold 8 (86.5%) while only 2 had homologs on Scaffold 12. Next, our synteny mapping of *AnoDis1.0* to the *A. carolinensis* assembly supported the identity of the *A. distichus* X chromosomes that was hypothesized based on prior cytogenetic work (Lisachov et al., 2019; Giovannotti et al., 2017; Kichigin et al., 2016): One neo-X chromosome should be syntenic to the ancestral X and fused with microchromosome 11. This neo-X chromosome was identified to be scaffold 8 (Fig. 3). The second neo-X chromosome, X_2_, should not be syntenic to the ancestral X chromosome, as it became sex-linked after the Y-autosome fusion (Lisachov et al., 2019). Supporting this, scaffold 12 was syntenic to the Y chromosome, comprised of scaffolds 13 and 16 as seen above, but not scaffold 8, (X_1_) (Fig. 4). Additionally, scaffold 12 (X_2_), was syntenic to the *A. sagrei* neo-X chromosome, which was formed as a result of a fusion between the ancestral X and ancestral chromosome 12 (Fig. 3) (Geneva et al., 2022).

Our depth of coverage, BLAST, and synteny results supported the hypothesized autosome-sex chromosome fusions previously predicted via cytogenetic and flow sorted chromosome sequencing analyses Lisachov et al. (2019); Giovannotti et al. (2017); Kichigin et al. (2016). Altogether, we identified scaffold 8 as X_1_, scaffold 12 as X_2_, and scaffolds 13 and 16 as parts of the neo-Y chromosome.

### Analyzing Evolutionary Pressures on Sex Chromosomes

To estimate whether neo-sex linked regions were undergoing degeneration and whether the neo-X chromosomes were experiencing the Fast-X effect, we compared the distributions of gene density, transposable elements, proportion of high and moderate effect SNPs, and K_a_/K_s_ values between the sex-linked scaffolds and similarly-sized autosomes (9, 10, and 11) in the assembly.

#### Evolutionary Pressure on the Y Chromosome

Our results support the prediction that Y-linked scaffolds have experienced degeneration relative to autosomes. First, we found that both of our candidate Y scaffolds (13 and 16) had fewer genes than similarly sized autosomes (W=916, p=7.163 *×* 10*^−^*^11^, onetailed, Wilcox signed-rank test). Second, although we found that 13 and 16 have an overall lower density of transposable elements than autosomes (W=943, p=1.445 *×* 10*^−^*^10^, one-tailed), relative to total scaffold length, these Y-linked scaffolds include regions with the highest densities of repetitive DNA. Additionally, Y-linked scaffolds have a greater proportion of high and moderate effect mutations (scaffolds 13 and 16 W=3803, p=5.246 *×* 10*^−^*^12^), and are under stronger selection than autosomes (scaffolds 13 and 16 W=60257, p=5.769 *×* 10*^−^*^12^) (Table 2).

**Table 2:** Median Values of Measured Sex Chromosome Attributes: Table showing the median values of each measured attribute for each sex-linked scaffold in comparison to similarly-sized autosomes.

| | Median Gene Density | Median Transposable Element Density | Median Proportion of High and Moderate Effect Variants | Median $K_a/K_s$ values |
| --- | --- | --- | --- | --- |
| Autosomes (scaffolds 9, 10, and 11) | 13 | 3427 | 0.41 | 1 |
| Scaffold 8 (neo-$X_1$) | 13 | 3181.5 | 0.73 | 4 |
| Scaffold 12 (neo-$X_2$) | 9.5 | 3565 | 0.73 | 3 |
| Scaffolds 13 and 16 (Y) | 5 | 2319 | 0.75 | 4 |

#### Evolutionary Pressures on X Chromosomes

Our results partially supported our predictions about degeneration and the Fast-X effect in both X_1_ (scaffold 8) and X_2_ (scaffold 12) chromosomes. Scaffold 12 had a lower gene density than autosomes (W=1656.5, p=0.02803), but Scaffold 8 did not (W=3356.5, p=0.3341). Contrary to our predictions, scaffold 8 had a lower density of transposable elements than autosomes (W=2158.5, p=0.002768, one-tailed), while transposable elements were not significantly more abundant on scaffold 12 than autosomes (W=2461, p=0.06792, one-tailed). However, Scaffold 12 had regions with particularly high densities of transposable elements, and in fact one region of Scaffold 12 encompasses the highest amounts of transposable elements across the genome, with more than 17,000 transposable elements in a single one-megabase window (Fig. 5., Table 2) With regard to selection strength metrics, both scaffolds 8 and 12 had a greater proportion of high and moderate impact SNPs than autosomes, suggesting that deleterious variants are accumulating on neo-sex chromosomes (scaffold 8 W=5202.5, p=2.873 *×* 10*^−^*^14^; scaffold 12 W=2744.5 , p=2.513 *×* 10*^−^*^9^). Finally, both scaffolds 8 and 12 are under stronger selection than autosomes, again supplementing our prediction that these neo-X chromosomes are experiencing the Fast X effect (scaffold 8 W=133538, p=8.741 *×* 10*^−^*^16^; scaffold 12 W=108716, (p=6.36 *×* 10*^−^*^14^) (Fig.5).

**Figure 5:**
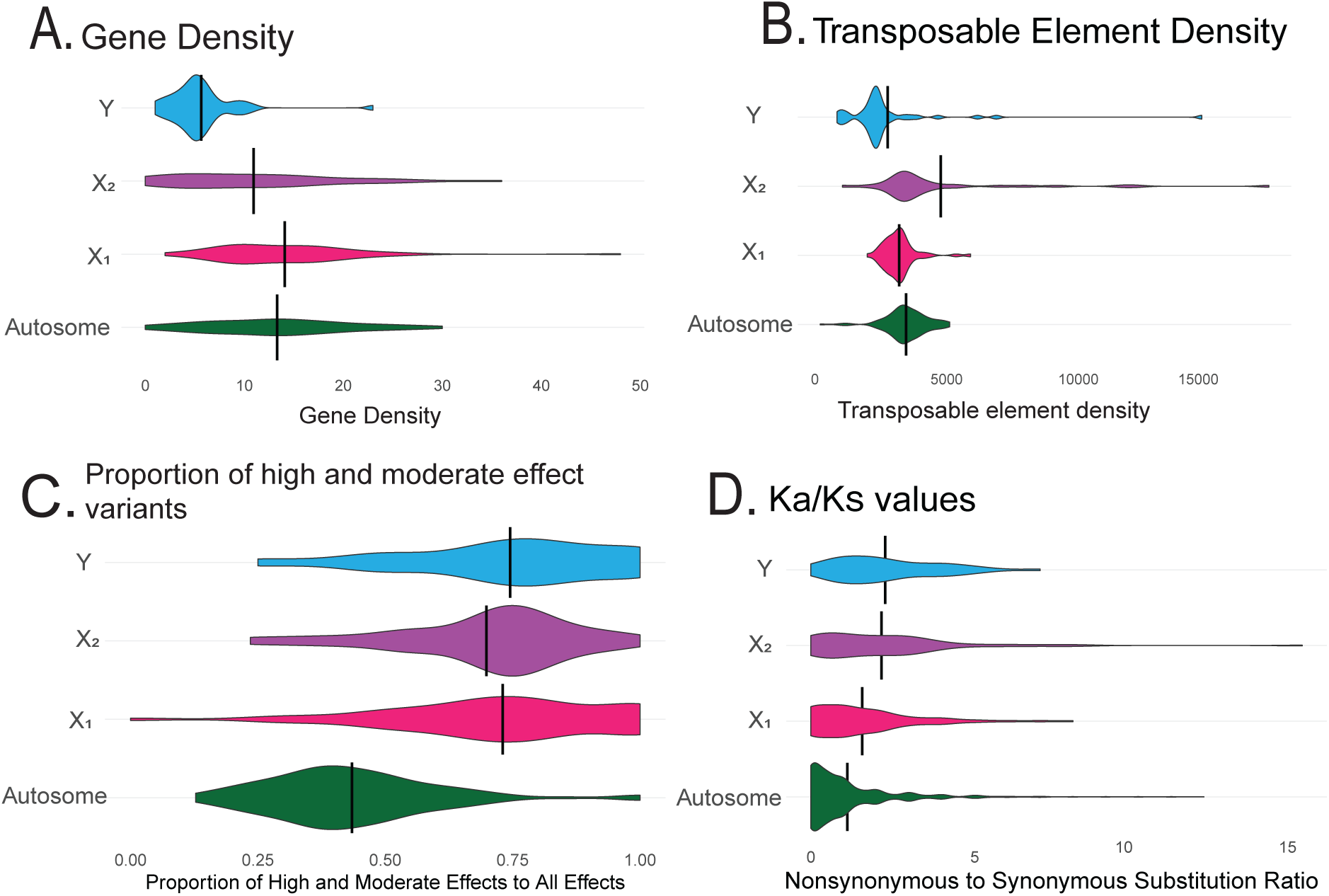
Violin plots comparing distributions of attributes of sex chromosomes: We compared different attributes of the X_1_, X_2_ and Y scaffolds to similarly-sized autosomal scaffolds (9, 10, and 11) within *AnoDis1.0*. The median of each distribution is indicated with a vertical line. **A.** Comparison of gene density across 1 megabase sliding windows, with the Y chromosome having the lowest gene density. **B.** Comparison of transposable element density across 1 megabase sliding windows, with both X_2_ and Y chromosomes experiencing an accumulation of transposable elements. **C.** Proportion of high and moderate effect variants across genes in each scaffold, with all sex chromosomes experiencing a higher proportion of these deleterious variants than autosomes. **D.** Density of synonymous to nonsynonymous mutation rates (K_a_/K_s_), a proxy for selection strength. All sex chromosomes have a higher ratio, highlighting that they may be under stronger selection pressures than similarly-sized autosomes.

### Nonrandom fusion of autosomes to sex chromosomes

We aligned the sex chromosomes of *A. distichus* to *A. sagrei*, a second *Anolis* species that has a neo-x chromosome, to test the prediction that certain autosomes within *Anolis* are more likely to become sex-linked (Fig. 3). Our data supported this prediction: Chromosome 12 in *A. carolinensis* (the hypothesized ancestral chromosomal arrangement), is an autosome, but in both *A. distichus* and *A. sagrei*, chromosome 12 is sex-linked. In *A. sagrei*, chromosome 12 fused with the ancestral X (Geneva et al., 2022), whereas in *A. distichus*, chromosome 12 fused with the ancestral Y 3.

## Discussion

We assembled and annotated a new, highly complete and contiguous genome for the Hispaniolan Bark Anole, *Anolis distichus*, to identify all chromosomes involved in the previously-predicted fusions and understand the evolutionary pressures on neo-sex chromosomes (Pennell et al., 2015; Lisachov et al., 2019; Gamble et al., 2014; Román-Palacios et al., 2018). This is, as far as we are aware, the first genome sequenced from a reptile with a multiple sex chromosome system (X_1_X_2_Y), providing valuable insight into the patterns of chromosomal fission and fusion driving the evolution of sex chromosomes (Pinto et al., 2023). We identified which autosomes were involved in the formation of the *A. distichus* neo-sex chromosomes via comparative synteny, finding support for the two previously predicted fusions that gave rise to the *A. distichus* X_1_X_2_Y sex chromosome system (Lisachov et al., 2019; Giovannotti et al., 2017; Kichigin et al., 2016). Next, neo-sex chromosomes can serve as a proxy to investigate the processes that gave rise to older, already degenerated sex chromosomes. However, it is unclear whether neo-sex chromosome degeneration follows predicted patterns associated with the classic model of sex chromosome degeneration, where regions of reduced recombination expand around a sex-determining locus (Abbott et al., 2017; Muller, 1964; Charlesworth et al., 2005). We assessed whether the neo-sex chromosomes in *A. distichus* matched theoretical predictions of how sex chromosomes should evolve, including whether they had undergone degeneration or had signatures of stronger selection strength, and found mixed results. Lastly, we add to a small but expanding number of studies that show patterns of the same autosome becoming sex-linked in multiple independent lineages (P̌senǐcka et al., 2024; Nozawa et al., 2021; O’Meally et al., 2012; Marshall Graves and Peichel, 2010; Sardell et al., 2021).

### Chromosomal Fusions in *A. distichus*

Previous research hypothesized that *A. distichus* experienced two fusions involving sex chromosomes and autosomes, resulting in their X_1_X_2_Y sex chromosome system; our *A. distichus* genome assembly provides another independent line of evidence supporting these fusions (Giovannotti et al., 2017; Lisachov et al., 2019; Kichigin et al., 2016). To start, we identified *AnoDis1.0* scaffolds 13 and 16 as two portions of the Y chromosome using synteny mapping, depth of coverage, and BLAST searches for genes present on the *A. carolinensis* Y chromosome. We hypothesized that, since the establishment of neo-sex chromosomes in the *Ctenonotus* clade of anoles is recent compared to the origin of the ancestral X and Y, neo-Y regions would still share syntenic regions to their neo-X counterparts. We performed pairwise synteny analyses between the *A. distichus* neo-sex-linked scaffolds, verifying that scaffold 8 and 12 are not syntenic to each other but are syntenic to portions of scaffolds 13 and 16. (Fig. 4). These two scaffolds cannot be assigned to neo-Y and ancestral-Y portions, as patterns of synteny between the neo-Y and neo-X regions depart significantly from co-linearity, suggesting the existence of multiple inversions and transversions between the ancestral and neo-Y portions of this chromosome (Supplementary Fig. 4), a pattern consistent with previous data from Y-linked regions in other taxa (Li et al., 2021; Xiao et al., 2020). However, we do observe elevated chromatin contact between the two scaffolds (13 and 16) hypothesized to encompass the *A. distichus* Y chromosome. Our inability to join scaffolds 13 and 16 together is most likely due to the Y chromosome’s regions of high repeat density, which makes chromosomal assembly challenging(Yamaguchi et al., 2021; Peona et al., 2021; Ezaz and Deakin, 2014).

We identified putative X_1_ (scaffold 8) and X_2_ (scaffold 12) chromosomes based on their depth of coverage, which were approximately half the depth of similarly-sized autosomes, and used BLAST search results to verify that scaffold 8 contained more than 86.5% of the known X-linked genes (Geneva et al., 2022) (Supplementary Fig. 3). Our synteny map between *A. distichus* and *A. carolinensis* confirms that *Anolis distichus* possesses two non-homologous X chromosomes (Fig. 3). The first, X_1_, is a consequence of fusions between the ancestral X and microchromosome 11. As such, X_1_ has an extensive region that is syntenic to the ancestral X, which is more than than 75 million years old and largely conserved across *Iguania* (Altmanová et al., 2018; Zheng and Wiens, 2016; Rovatsos et al., 2014a) The second, X_2_, is syntenic to microchromosome 12, formed as a result of the Y-autosome fusion (Fig. 1) . Our results support the existence of the two sex chromosome and autosome fusions that *A. distichus* has undergone to ultimately form its X_1_X_2_Y chromosome system, and our identification of the chromosomes involved in these fusions is identical to what was predicted by previous cytogenetic work (Lisachov et al., 2019; Kichigin et al., 2016; Giovannotti et al., 2017).

### Degeneration in the Y Chromosome

As predicted, we found that *AnoDis1.0* Y scaffolds had the lowest gene density of all chromosomes studied, including similarly-sized autosomes and the X_1_ and X_2_ chromosomes (Fig. 5, Table 2). While the overall distribution of TEs on the Y scaffolds was lower than autosomes, there were regions with extremely high TE density (Fig. 5), consistent with the expected non-random distribution of TEs throughout the genome (Lawlor and Ellison, 2023; Hollister and Gaut, 2009). Overall, these findings are consistent with the classic theory of sex chromosome degeneration, and empirical data from other taxa, where the Y chromosome undergoes degeneration that results in lower gene densities and an accumulation of transposable elements (Ahmad et al., 2020; Charlesworth and Charlesworth, 2000; Sakamoto and Innan, 2021; Śliwińska et al., 2016).

### Degeneration and Selection in the neo-X Chromosomes

We tested whether formerly autosomal regions of the neo-X-linked chromosomes of *A. distichus* show patterns of TE and gene density consistent with recombination suppression and the early establishment of evolutionary strata (Wright et al., 2016; Palmer et al., 2019; Charlesworth, 2021). As expected under a complex scenario of multiple and independent fusions to X and Y chromosomes, patterns differed between X_1_ and X_2_: X_1_ had no significant difference in gene density with respect to autosomes, but did have significantly more transposable elements (TEs). Conversely, X_2_ had a lower gene density when compared to autosomes. Although there was no significant difference when compared to autosomes, X_2_ also had regions with extremely high TE density and has the highest density of transposable elements in a window across all scaffolds (Fig. 5, Supplementary Fig. 6). Again, an uneven distribution of TEs is expected, given that distinct TEs accumulate non-randomly across the genome (Lawlor and Ellison, 2023; Hollister and Gaut, 2009; Langmüller et al., 2023). These patterns indicate that X_2_ has accumulated transposable elements within specific regions, perhaps as a consequence of a reduced recombination rate following the Y-autosome fusion.

In a scenario of degeneration, we would expect some degree of pseudogenization and degeneration as recombination ceases between X_2_ and the Y (Mrnjavac et al., 2023; Nozawa et al., 2016, 2021). We have evidence for this process in the reduction in gene density for X_2_. We cannot, however, rule out that a lower gene density is an an inherent property of the ancient microchromosome 12, since this ancient microchromosome has also become sex-linked in *A. sagrei*. Its assembly in *A. carolinensis* is highly fragmentary, with the assignment of scaffolds to microchromosome 12 in *A. carolinensis* still relying on the alignment of short reads from flow sorted chromosomes to the *A. carolinensis* genome (Kichigin et al., 2016; Giovannotti et al., 2017). Nevertheless, our results suggest that patterns of degeneration for neo-sex-linked chromosomes depend not only on time since fusion, but also on whether linkage has occurred to the sex-limited (Y) or non-sex-limited chromosome.

We hypothesized that sex chromosomes would be under different selection pressures than autosomes. More specifically, we predicted that both neo-X chromosomes would be experiencing the Fast-X effect, where X chromosomes experience faster rates of evolution as recessive beneficial mutations are exposed in the heterogametic sex, but recessive negative mutations are hidden in the homogametic sex, leading to an overall more “efficient” process of evolution (Charlesworth et al., 1987; Meisel and Connallon, 2013; Darolti et al., 2023). Taken together, our SNP-Eff and K_a_/K_s_ results showed that genes on sex chromosomes are under recurrent positive selection, likely in part due to the Fast-X effect (Fig. 5, Table 2) (Yang and Bielawski, 2000; Li, 1993). The Fast-X effect can be observed quickly after sex chromosome turnover occurs, so it is not surprising that we observe X_1_ and X_2_ to be experiencing this (Darolti et al., 2023). Our result aligns with a previous study in *A. carolinensis* (Rupp et al., 2016). Overall, the trends seen when comparing X_1_ and X_2_ to autosomes indicate that the patterns in X_1_ align with expected theory for X chromosomes, and highlight that X_2_ is now evolving as a sex chromosome rather than an autosome after the fusion event.

### Independent transitions from autosomes to sex chromosomes

We predicted that sex chromosome and autosome fusions are not random, and that the same autosome may be repeatedly involved in fusions to create neo-sex chromosomes (Sardell et al., 2021; O’Meally et al., 2012; Marshall Graves and Peichel, 2010). Our results, along with the recently assembled *A. sagrei* genome, indicate that the ancestral microchromosome 12 has become sex-linked in at least two highly diverse anole clades: *Ctenonotus* and *Norops* (Fig. 3) (Geneva et al., 2022). Phylogenetic relationships among anole subgenera at the base of the *Anolis* phylogeny are notoriously difficult to resolve, however two recent studies support the idea that *Norops* and *Ctenonotus* are sister clades (Alföldi et al., 2011; Mahler et al., 2016). Therefore, we propose three evolutionary scenarios for microchromosome 12’s neo-sex linkage: (i) The ancient microchromosome 12 fused to the Y-chromosome in the ancestor of the *Norops* + *Ctenonotus* clade, undergoing a secondary fusion to the ancestral X of *Norops*, but not in the ancestor of *Ctenonotus*; (ii) The ancient microchromosome 12 and the Y fused independently the ancestors of *Norops* and *Ctenonotus* clades, while a fusion between microchromosome 12 and the ancient X occurred exclusively in the *Norops* clade - either prior to, or after the Y-fusion; and (iii) The ancient X and ancient Y fused to the autosomal homologs of microchrosome 12 in the common ancestor of *Ctenonotus* and *Norops*, followed by a fission of the Neo-X exclusively in the *Ctenonotus* clade.

We favor hypothesis (i), where the same microchromosome has fused independently more than once within a clade. Indeed, independent autosome to sex-chromosome fusions across two closely-related lineages have been documented in a diversity of taxa: In flies, for example, a region of the genome known as Muller Element C independently became part of the neo-Y chromosomes in *D. miranda* and *D. albomicans*, and genes present on these chromosomes were found to be undergoing patterns of parallel degeneration (Nozawa et al., 2021). In stickleback fish, chromosome LG12 became sex-linked in two different lineages, once through a fusion with the Y chromosome in *G. wheatlandi*, and once in *P. pungitus*, potentially through introgression (Kitano et al., 2009; Sardell et al., 2021; Dixon et al., 2019). In snakes, homology of chromosome content across a diversity of sex chromosome systems suggest that certain regions of the genome may be more likely to become sex-linked (P̌senǐcka et al., 2024). Our results support that certain autosomes may be repeatedly recruited to become to sex chromosomes (reviewed in (O’Meally et al., 2012; Marshall Graves and Peichel, 2010)). However, synteny alone cannot reject any of the aforementioned hypotheses, and thus highlight the need for continued sampling of taxa in *Anolis*, where multiple sex-determining systems have evolved independently (Gamble et al., 2014).

The repeated recruitment of the same chromosome as part of a neo-sex chromosome system also facilitates future comparative studies on the evolution of dosage compensation, a process in which necessary gene products are equalized between the heterogametic and homogametic sex (Charlesworth, 1998; Gu and Walters, 2017). Dosage compensation is a key player in sex chromosome evolution, and potentially adaptation and speciation (Engelstädter, 2008; Gu and Walters, 2017). In *A. carolinensis*, dosage compensation causes X-linked genes to be up-regulated in males, equalizing gene product (Rupp et al., 2016; Marin et al., 2017). Future work should test whether X up-regulation occurs in other species of *Anolis*, and if so, how this pattern is established in systems with neo-sex chromosomes such as *A. distichus*.

Lastly, the Hispaniolan Bark anole is a highly polymorphic anole species, and shows a rare polymorphism of its dewlap, a key secondary sex trait. Across most anole species, dewlap color patterns are fixed within species, often being hypothesized to be part of a classic lock-key system of species identification (Fitch and Hillis, 1984; Sigmund, 1983; Vanhooydonck et al., 2009). Within *A. distichus*, however, dewlap color varies with environment, being hypothesized to be driven by local selection for signal optimization (Behere et al., 2024; De Mello et al., 2021). Furthermore, *A. distichus* has also been shown to include multiple phenotypically distinct populations (classified as subspecies) that come in contact independently across the island of Hispaniola, showing different levels of reproductive isolation at each contact zone (Geneva et al., 2015; MacGuigan et al., 2017; Glor and Laport, 2012; Ng et al., 2016; Case and Williams, 1984; Ng and Glor, 2011). Therefore, the *A. distichus* genome will also allow future microevolutionary studies on the processes underlying adaptation and speciation within anoles, including the roles of neo-sex chromosomes and genomic architecture in these processes.

Our highly complete and contiguous genome assembly of the Hispaniolan Bark anole (*A. distichus*) is, to our knowledge, the first assembly of a reptile species with a multiple sex chromosome system. *Anolis distichus* is part of *Ctenonotus*, a unique clade within *Anolis* that has an X_1_X_2_Y system originating from independent autosome-to-sex-chromosome fusions to both the X and Y chromosomes. Our syntenic analyses not only identify which ancestral autosomes fused to sex chromosomes, but also identify the regions within these chromosomes where depth of coverage, heterozygosity, and other metrics associated with the suppression of recombination are expected to change. It will be interesting to see if and how these patterns of degeneration vary both within and between species of anoles within the *Ctenonotus* clade, and whether they correlate with patterns of diversification of these species across the anole adaptive radiation. Altogether, our assembly adds to a growing library of genomes of anoles, which are considered one of the premier model systems for the study of ecology and evolutionary biology, enabling future testing of evolutionary hypotheses through comparative approaches on a genomic level that will further our understanding of sex chromosome evolution and diversification as a whole.

## Acknowledgments

We thank Brewster Kingham, Dr. Erin Bernberg, Dr. Olga Shevchenko, and Mark Shaw at the University of Delaware Sequencing and Genotyping Center for PacBio extraction and sequencing. We thank Dovetail Genomics for Omni-C library preparation, ACU staff at KU for lizard care, KUMC GSC staff for library preparation and sequencing, and Rutgers OARC, XSEDE, and the KU CRC for computational resources.

## Data Availability Statement

Data are available at NCBI under BioProject PRJNA1200026, BioSample SAMN45894292. This data is currently embargoed but will be released upon publication. Annotation, repetitive elements, SNP-calling files, and chromatin contact files are similarly embargoed but will be available upon publication at the Harvard Dataverse: https://doi.org/10.7910/DVN/HJD8VT.

Scripts created to perform the analysis can be found at: https://github.com/chfal/distichus.

## Supplemental Figures

**Supplemental Figure 1:**
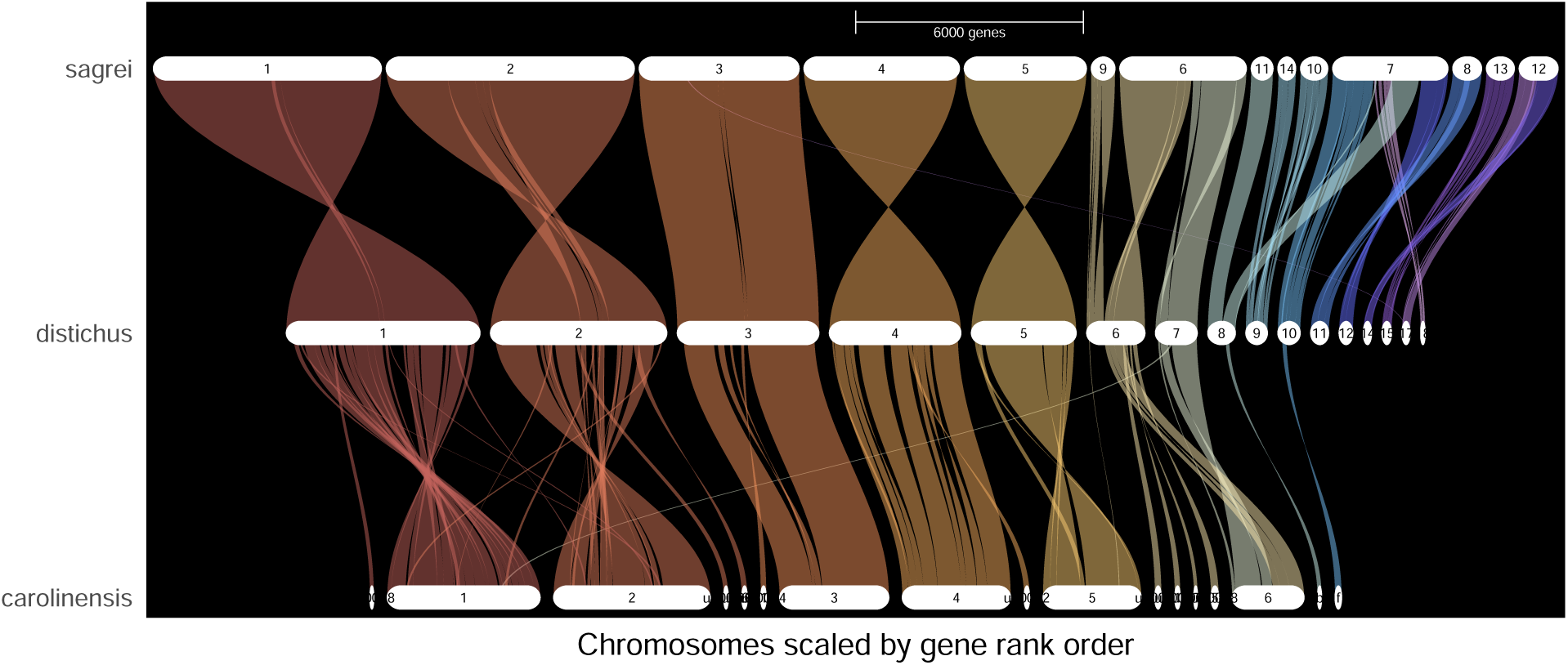
Macrochromosome synteny: Genespace syntenic relationships of macrochromosomes in *A. carolinensis* compared to *A. sagrei* and *A. distichus*.

**Supplemental Figure 2:**
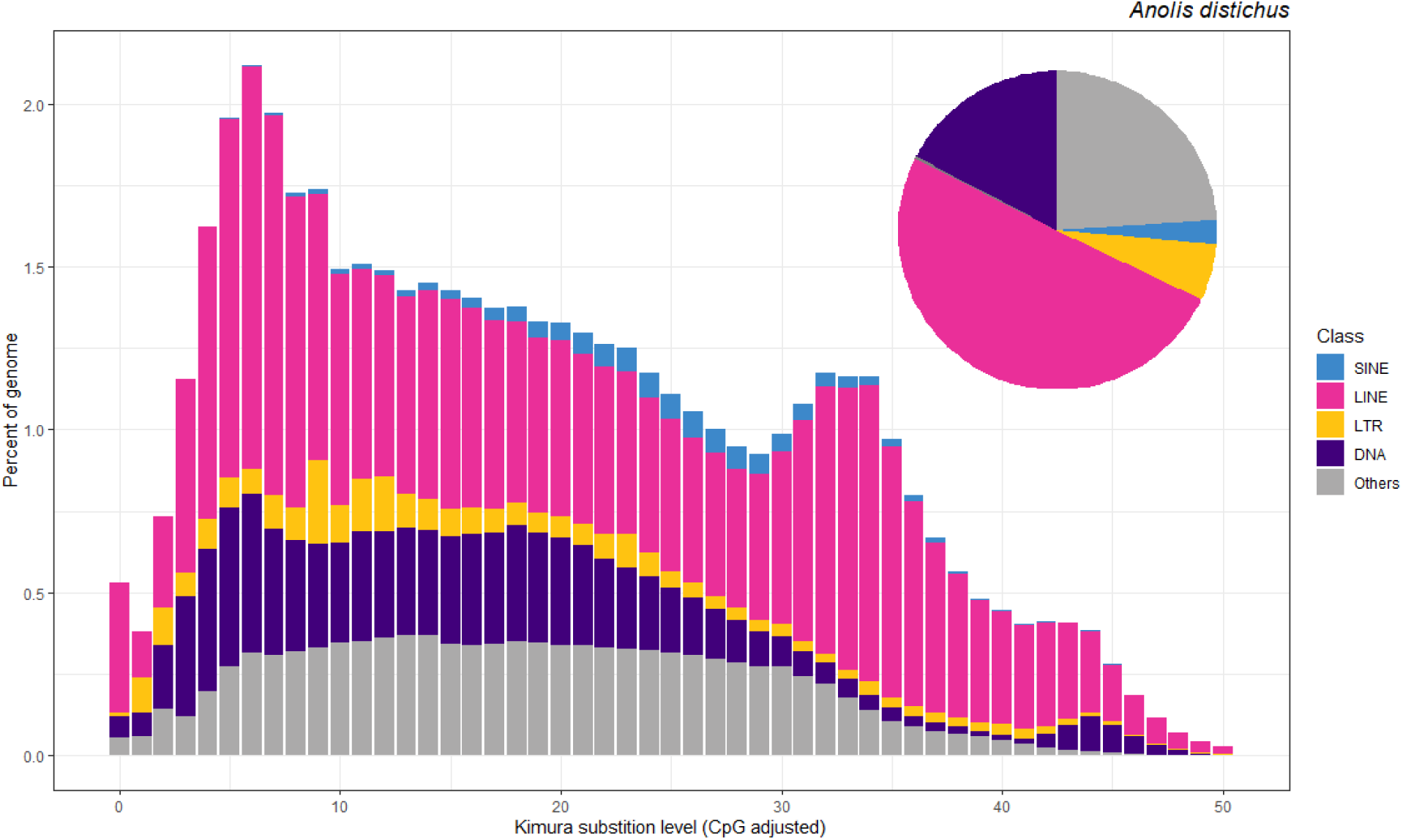
Repeat Landscape: Repeat landscape of *AnoDis1.0* assembly with colors representing categorized, identified repeat elements.

**Supplemental Figure 3:**
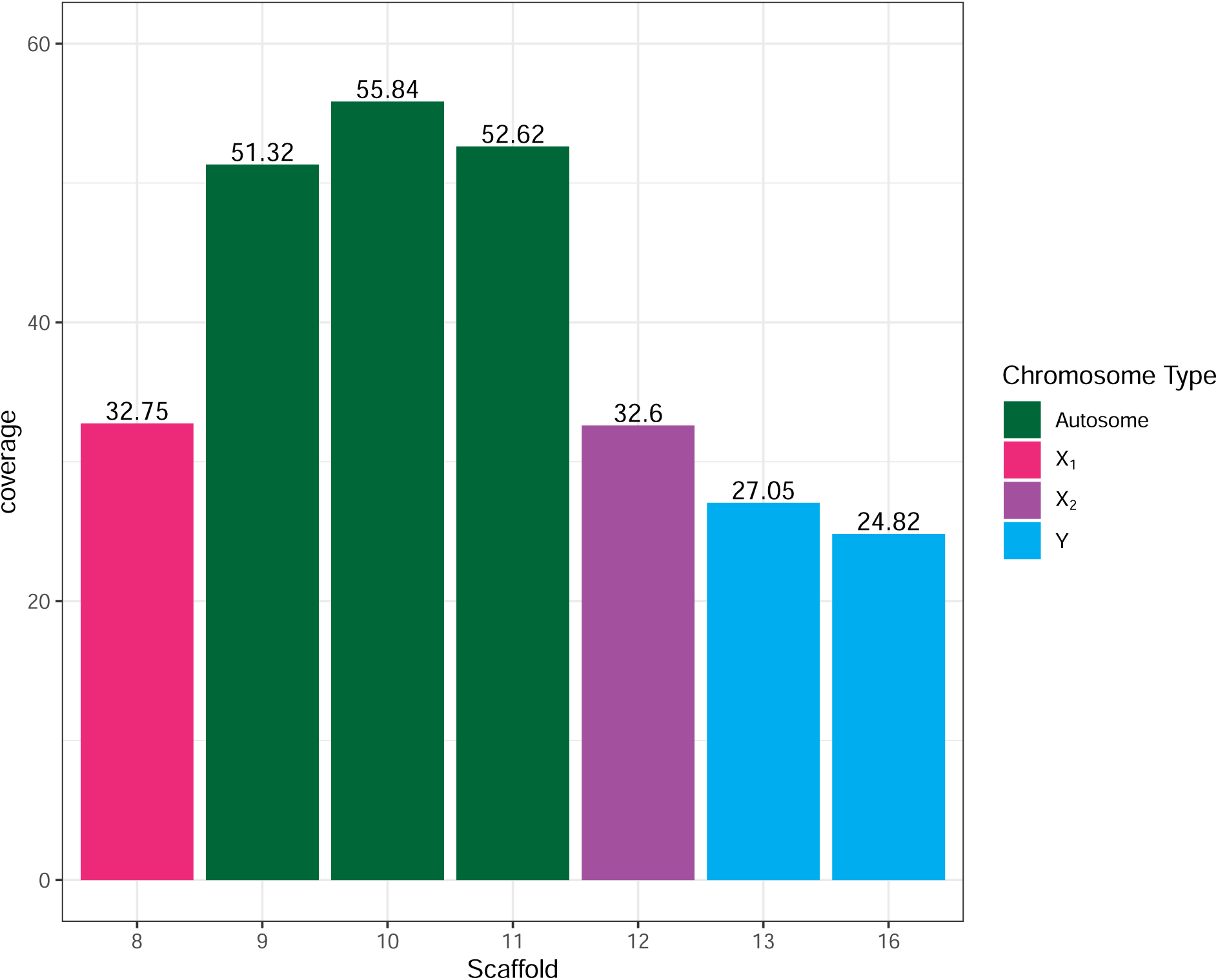
Coverage depth by scaffold: Histogram of coverage by scaffold between autosomes and identified sex-linked scaffolds. The coverage depth of putative sex-linked scaffolds is much lower than similarly-sized autosomes, which is expected since the genome animal is a male and therefore should be hemizygous for X_1_, X_2_, and Y.

**Supplemental Figure 4:**
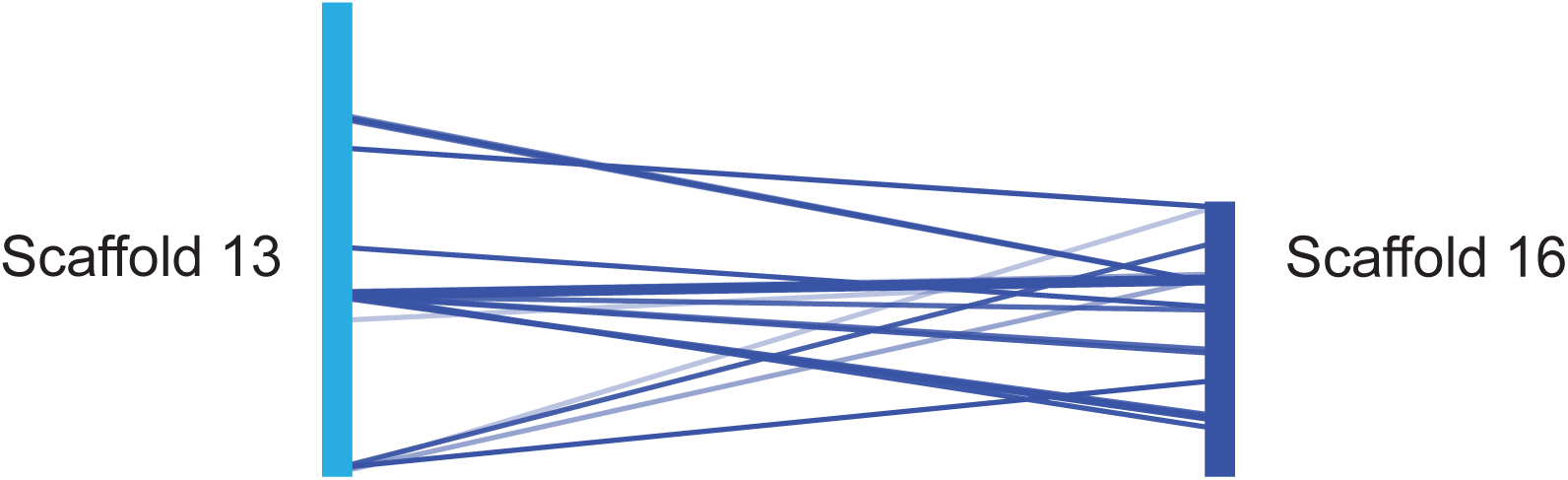
Synteny between Y scaffolds: Syntenic mapping of scaffold 13 to scaffold 16 in *AnoDis1.0* shows the linkage between the two portions of the Y chromosome in our assembly. The Y chromosome is assembled in two pieces likely due to the challenging nature of assembling highly repetitive chromosomes.

**Supplemental Figure 5:**
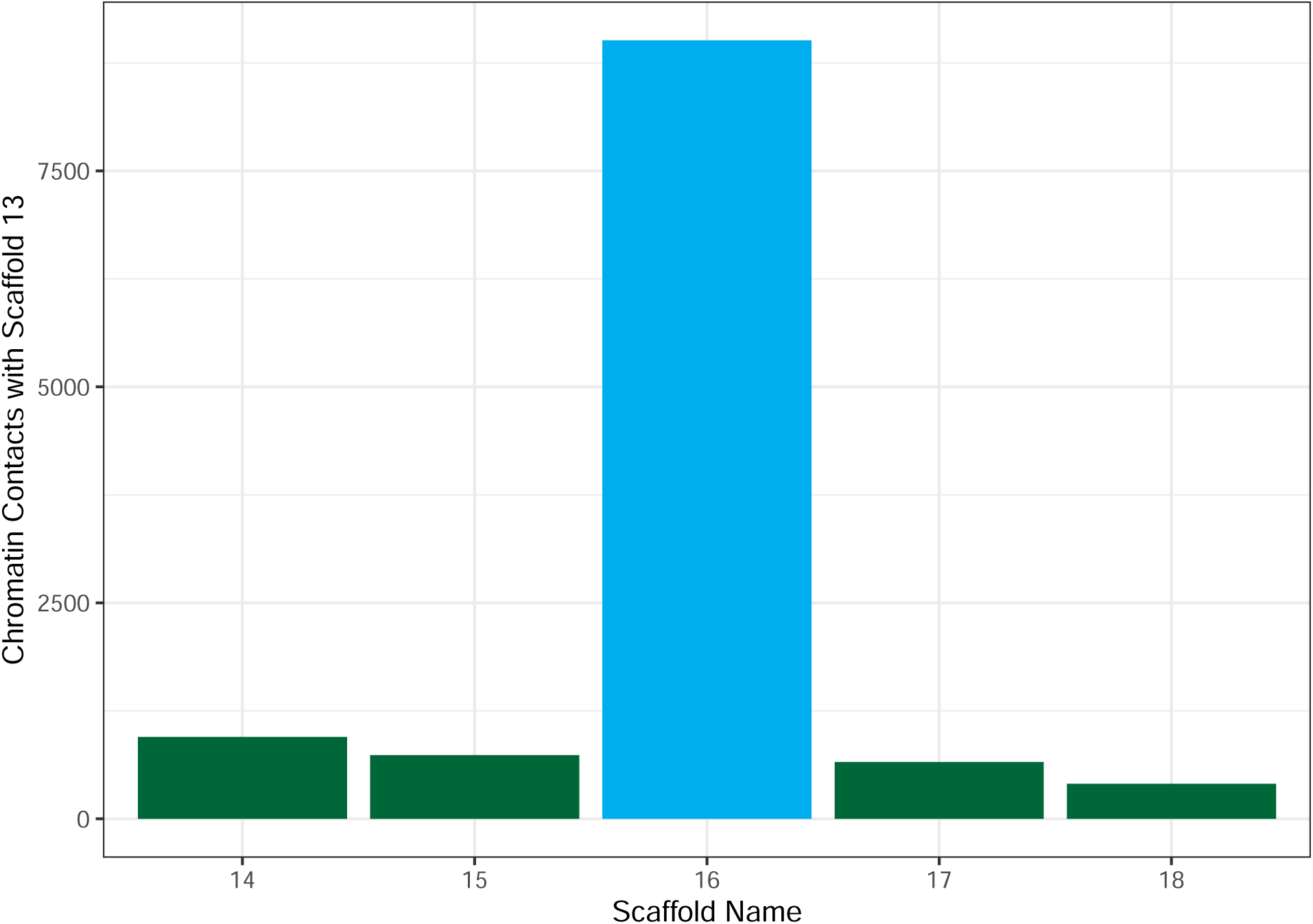
Chromatin Contact Between Scaffold 13 and Other Scaffolds: Counts of chromatin contacts between scaffold 13 and other similarly-sized scaffolds (14, 15, 16, 17, and 18). There is an extremely large number of contacts between scaffold 13 and scaffold 16 as compared to the other scaffolds, supporting the idea that the *Anolis distichus* Y chromosome was assembled in two pieces.

**Supplemental Figure 6:**
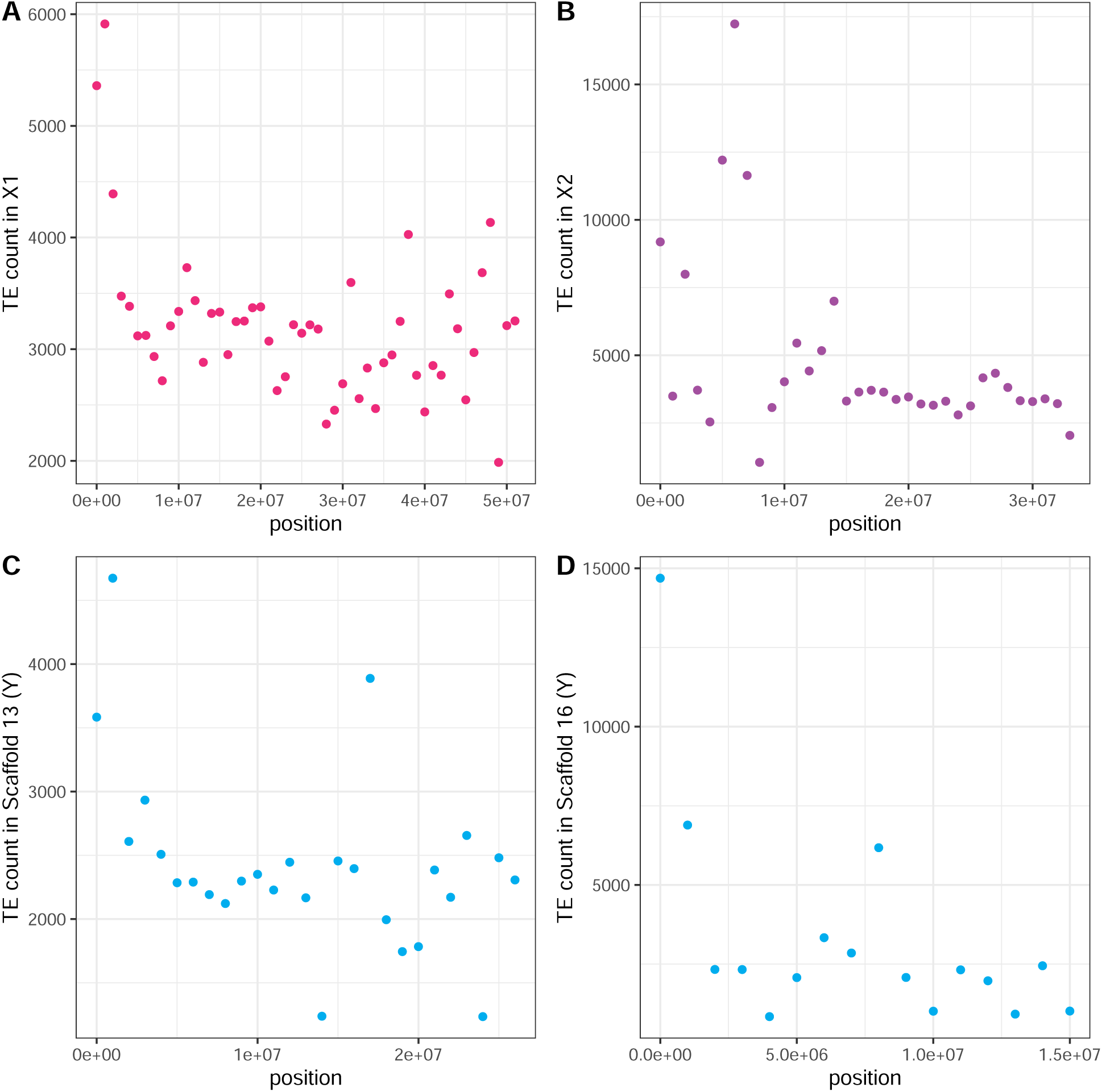
Transposable element counts across sex-linked scaffolds: Transposable element count across 1 MB windows in all four sex-linked scaffolds; X_1_ (A), X_2_ (B), and the Y (scaffold 13, (C); scaffold 16, (D). X_2_ and Y chromosomes have among the highest numbers of transposable elements.

